# Fine-scale movement data reveal surface foraging and nocturnal flight activity in the endangered Bermuda petrel

**DOI:** 10.1101/2024.05.08.593164

**Authors:** Paolo Becciu, Allison Patterson, Carina Gjerdrum, Jeremy Madeiros, Letizia Campioni

**Author notes:** Corresponding author, Letizia Campioni.

## Abstract

Foraging behavior plays a fundamental role in animal fitness and population dynamics, particularly in marine ecosystems where seabirds like petrels (small Procellariiformes) showcase a diverse array of foraging strategies finely adapted to the pelagic environment. The extent and remote nature of their foraging grounds makes direct observation of foraging behaviour impractical, thereby requiring the use of remote tracking technologies. We deployed miniaturized multi-sensor biologgers and collected fine scale movement data to investigate the at-sea behaviours of the Bermuda petrel *Pterodroma cahow*, a poorly studied and highly threatened gadfly petrel, specialised on mesopelagic prey. GPS-tracking data revealed extensive foraging trips, in consistent directions, over remote oceanic regions. Time-depth-recorders provided new insights into petrel feeding techniques suggesting that the meso-bathypelagic prey targeted by petrels must be available in the very upper layer of the water surface, given their very limited diving activity (99.99% of dives had 0.1 m of depth). Accelerometer data revealed 3 flying- and 3 water-related behaviours. Flying behaviours reflected the expected dynamic soaring flight strategy of procellariforms; individuals spent more than three-quarters of their time in flight with flying-non-flapping being the most common behaviour under all conditions. The behaviour classified as “flying-intensive” was infrequently observed but could indicate aerial dipping, a characteristic foraging technique of *Pterodroma* species. The remaining time was spent in three water behaviours: active, inactive and intensive, with the latter being less common but thought to reflect scavenging and prey seizing. Flying-related behaviours increased with negative sun elevation values, highlighting greater flight activity during night compared to the day, while water behaviours were more common during the day. While some of our findings may require further validation to confirm their relevance to foraging behaviour, our work offers new and valuable insights to consider when assessing the extent and nature of offshore anthropogenic-related risks faced by petrels.

## Introduction

Foraging is a crucial component of animal behaviour with direct implications for individual fitness, survival and population demography [1]. In marine ecosystems, foraging behaviour establishes a predator–prey link influencing both the trophic connectivity and the carbon flux in the oceans [2, 3]. Identifying a predator’s at-sea behaviour and foraging strategy is beneficial for understanding the species’ ecological requirements and roles in ecosystems. Procellariform seabirds such as petrels and albatrosses are upper-level predators of pelagic food webs showing an impressive range of foraging strategies and feeding techniques [4]. Many of them are well adapted to perform long-distance dynamic soaring flight with minimal energy expenditure [5], enabling them to spend a significant portion of their life in the open ocean while in search of food resources.

Advances in tracking technology have made it possible to study the oceanic movements of many pelagic predators, including seabirds, thus unveiling different aspects of their foraging behaviour and movements [6, 7] alongside the biophysical conditions they encounter at sea [8]. Despite this, aspects of flight behaviour, activity patterns and feeding techniques, particularly of smaller seabirds such as petrels, remain poorly understood [9]. Until recently, the feeding behaviour of petrels was derived primarily from analysing stomach contents and regurgitates [10–12], and although this method has contributed to a better understanding of the ecological role of these small predators in food webs, the addition of behavioural data at sea has raised new questions. For example, how do petrels gain access to their mesopelagic prey species that ascend from depth to the water surface after sunset [13] (i.e. diel vertical migration) when tracking devices have shown petrels are primarily foraging during the daytime [14].

Gadfly petrels of the genus *Pterodroma* are pelagic seabirds morphologically and anatomically adapted to exploit wind energy to perform prolonged and efficient dynamic soaring flights [15], during which they mostly glide on the ocean surface with limited flapping-flight [16]. The species within this group are also ecologically very similar, with a diet based predominantly on deep sea mesopelagic prey [17, 18], which they capture using different feeding techniques [19]. Indeed, *Pterodroma* petrels are believed to forage while in flight by contact “dipping” to the surface (also known as “stooping” or “aerial dipping,” that is, taking up prey in flight while barely touching the sea surface; [20]), which is considered a shared trait among congeners [21]. They may also feed while sitting on the water through surface-seizing or scavenging, and less frequently by searching for prey underwater [22, 23]. Due to their specialization on mesopelagic prey, including fish (mostly Myctophids) and cephalopods [17], which in dark conditions migrate vertically through the water column [13], petrels are expected to increase their foraging activity at night [12, 17, 19]. *Pterodroma* petrels are not regarded as proficient divers unlike shearwaters or diving-petrels [16, 20, 23–27]. This interspecific variation is based on the analysis of morphological traits (e.g. body size, long wings, lack of laterally compressed legs) and physiological characteristics; however, the diving capacity and performance in this genus are still anecdotal or poorly investigated.

We studied the at-sea behaviour of the Bermuda petrel (*Pterodroma cahow*), a globally endangered gadfly petrel endemic to the western North Atlantic that breeds exclusively in the Bermuda Islands [28, 29]. To date, only two studies have focused on the foraging ecology of this species [18, 30]. Campioni et al. [18] showed that the petrel’s flight, similar to other congeners, can be modulated through wind selection, likely facilitating extended and prolonged foraging trips while minimizing energy expenditure. Although the diet of the Bermuda petrel is dominated by meso- and bathypelagic fish (mostly Myctophids) and cephalopods [18], which would indicate substantial nocturnal foraging activity, a previous analysis of GPS data suggested an equivalent amount of “searching” behaviour (i.e. putative foraging) vs “transit” by both day and night [18]. In order to improve our understanding of the species’ daily activity budget, we combined the use of GPS, tri-axial accelerometer, and time-pressure sensors to collect complementary and high-resolution information to (i) characterise petrel foraging areas, (ii) identify behavioural modes when they are in flight and on the water including diving activity, and (iii) test whether these behaviours change in relation to the diel cycle.

## Materials and Methods

### Biologging data collection

Biologging devices were deployed on 25 incubating Bermuda petrels at Nonsuch Island, Bermuda (32.347°N, 64.663°N) between 22 January and 9 February 2023. Birds were temporarily equipped with loggers on their backs or tails using Tesa tape 4651. Most loggers (19 out of 25) were recovered 13–26 days later. Three logger types were used in this study: (1) Axy5 loggers (Technosmart, Italy, 2.5 g) that included triaxial accelerometers (25 Hz), tri-axial magnetometers (1 Hz), and time-pressure sensors (1 Hz) were deployed on the backs of 9 petrels; (2) AxyTrek loggers (Technosmart, Italy, 5 g) that included GPS sensors (1 fix every 5 min), tri-axial accelerometers (25 Hz), and time-pressure sensors (1 Hz) were deployed on the backs of 5 petrels; and (3) Nano-fix GEO loggers (PathTrack, United Kingdom, 3.4 g) that only recorded GPS locations (1 fix every hour) were deployed on the tail of 11 petrels (for details on attachment method see Campioni et al. [18]). Devices were deployed on breeding adults that weighed between 300 and 385 g (N=25); thus, the heavier loggers (AxyTrek) corresponded to 1.3–1.7% of body mass while the lighter loggers (Axy5) corresponded to 0.8–0.65% of body mass. Data from deployments and biologgers are included in Table 1.

**Table 1:** Summary of tracking data and foraging trips of Bermuda petrels tracked from Nonsuch Island in 2023.

| Total Track |  |  |  |  |  |  |  | Foraging Trip |  |  |  |  |  |
| --- | --- | --- | --- | --- | --- | --- | --- | --- | --- | --- | --- | --- | --- |
| Colony | Tag type | Nest | Bird | Sex | Start | End | Duration (days) | Trip | Start | End | Duration (days) | Maximum distance (km) | Notes |
| A | Axy5 | R819 | E0368 | M | 2023-01-25 20:27 | 2023-02-03 1:03 | 8.2 | 1 | 2023-01-26 9:20 | 2023-01-31 1:35 | 4.7 | NA | Complete |
|  | PathTrack | R820 | E0487 | F | 2023-02-02 2:07 | 2023-02-17 18:07 | 15.7 | 1 | 2023-02-08 2:07 | 2023-02-17 6:07 | 9.2 | 1493 | Complete |
|  | PathTrack |  | E0243 | M | 2023-01-23 2:03 | 2023-02-08 3:04 | 16.0 | 1 | 2023-01-26 11:03 | 2023-02-07 22:04 | 12.5 | 1503 | Complete |
|  | Axy5 | R821 | E0362 | M | 2023-01-23 0:00 | 2023-01-31 3:28 | 8.1 | 1 | 2023-01-24 6:20 | 2023-01-27 3:35 | 2.9 | NA | Complete |
|  | PathTrack | R834 | E0161 | F | 2023-01-29 2:00 | 2023-02-12 19:00 | 14.7 | 1 | 2023-02-01 6:00 | 2023-02-11 2:00 | 9.8 | 1100 | Complete |
|  | Axy5 |  | E0182 | M | 2023-01-25 21:07 | 2023-02-03 5:37 | 8.4 |  |  |  |  | NA | In burrow |
|  | AxyTrek | R835 | E0220 | M | 2023-01-26 0:00 | 2023-02-02 8:03 | 7.3 | 1 | 2023-02-01 7:35 | 2023-02-02 8:05 | 1.0 | [549] | Incomplete |
|  | Axy5 |  | E0401 | F | 2023-01-25 20:15 | 2023-02-03 3:52 | 8.3 |  |  |  |  | NA | In burrow |
|  | AxyTrek | R836 | E0171 | M | 2023-02-09 0:00 | 2023-02-18 22:01 | 9.9 | 1 | 2023-02-18 2:55 | 2023-02-18 22:00 | 0.8 | [1075] | Incomplete |
|  | PathTrack | R837 | E0801 | F | 2023-01-29 2:02 | 2023-04-01 14:06 | 62.5 | 1 | 2023-01-30 8:02 | 2023-02-01 6:02 | 1.9 | 232 | Complete |
|  |  |  |  |  |  |  |  | 2 | 2023-02-11 7:02 | 2023-02-23 23:05 | 12.7 | 1433 | Complete |
|  | PathTrack | R839 | E0363 | F | 2023-01-26 2:03 | 2023-02-19 19:05 | 24.7 | 1 | 2023-02-04 5:02 | 2023-02-18 2:05 | 13.9 | 1003 | Complete |
|  | PathTrack | R840 | E0484 | M | 2023-01-26 2:02 | 2023-02-08 3:02 | 13.0 | 1 | 2023-01-28 5:01 | 2023-02-03 3:02 | 5.9 | 675 | Complete |
| B | AxyTrek | B12 | C0888 | F | 2023-02-10 0:00 | 2023-02-18 7:33 | 8.3 | 1 | 2023-02-18 1:55 | 2023-02-18 7:35 | 0.2 | [231] | Incomplete |
|  | Axy5 |  | E0252 | M | 2023-01-26 21:10 | 2023-02-04 4:47 | 8.3 |  |  |  |  | NA | In burrow |
|  | Axy5 | B2 | C0901 | M | 2023-01-26 20:00 | 2023-02-04 4:04 | 8.3 | 1 | 2023-01-27 6:35 | 2023-02-03 4:05 | 6.9 | NA | Complete |
|  | Axy5 | B8 | E0083 | F | 2023-02-10 0:00 | 2023-02-18 2:18 | 8.1 | 1 | 2023-02-10 0:00 | 2023-02-18 2:20 | 8.1 | NA | Incomplete |
|  | Axy5 |  | C1036 | M | 2023-01-26 21:00 | 2023-02-03 17:59 | 7.9 | 1 | 2023-01-28 7:15 | 2023-02-03 18:00 | 6.4 | NA | Incomplete |
|  | PathTrack | B9 | E0552 | F | 2023-01-27 2:00 | 2023-02-14 23:00 | 18.9 | 1 | 2023-02-07 1:00 | 2023-02-14 23:00 | 7.9 | 1241 | Complete |

### Defining foraging trips and at-sea distribution

Foraging trips were defined as any period between a bird’s departure and subsequent return to the colony, excluding locations within a range of 1 km from the colony (thus removing positions on land and those possibly related to rafting in the proximity of the colony [31]). Only tracks with at least one complete foraging trip were used to calculate the maximum distance from the colony and the overall temporal duration of the trip. We fitted discrete-time Hidden Markov models (HMMs) to the GPS data using the package moveHMM [32] in R [33] using 2-state HMMs to classify the behavioural states based on the distance travelled (step length) and the change of movement direction (turning angle), similarly to [18]. The most likely sequence of behavioural states was inferred using the Viterbi algorithm [34]. We assumed that, along the tracks, the birds were in one of the following behavioural states: ‘transit’, in which birds were moving at high speed at a persistent heading; or ‘search’, in which birds were performing convoluted movements at a comparatively lower speed [35, 36]. We considered the search state to be associated with food searching or putative foraging behaviour, although it can also include resting behaviour (for more details see Campioni et al. [18]). Next, we fitted a continuous-time movement model to calculate a population-level utilization distribution (UD) of foraging areas using 95%, 75%, 50% and 25% autocorrelated kernel density estimate (AKDE) for all the individuals using the functions `akde` and `pkde` from the package `ctmm` [37, 38]. We fitted the AKDE using all tracks but only those locations labelled as ‘searching’ by the HMM and excluding locations within 50 km of the colony (see Campioni et al. [18]).

### Diving Activity

Depth data were collected at 1 Hz with a resolution of 0.005 m. Two outliers with depths greater than 15 m were removed from the dataset (<0.00003% of all depth measurements) and these missing values were replaced with linearly interpolated values. These extreme observations were treated as depth sensor anomalies because they were isolated events that would have required travelling at vertical speeds >15 m/s. A zero-offset correction (ZOC) was used to correct for temporal variation in the accuracy of the depth sensor measurements [39]. We calculated the ZOC as the 10th percentile of depth over a 10-minute moving window. ZOCs ranged from 0– 0.49 m. The ZOC was then subtracted from all depth measurements and corrected values <0 m were fixed at 0 m. We examined histograms of depth measurements while birds were engaged in foraging trips (see below for details of behavioural classification) for evidence of diving activity.

### Accelerometer behavioural classification

The wide-ranging behaviour of *Pterodroma* petrels makes it difficult to quantify detailed behavioural states at-sea. Triaxial accelerometers are a tool that can be used to collect high-resolution activity data from wildlife over long periods of time even when individuals are not readily observable [40–44]. Accelerometers record changes in acceleration on three planes: surge acceleration (forward-backward), sway (left-right), and heave (up-down). Behavioural patterns can often be readily identified by an observer looking at plots of different measures of acceleration. However, for individuals tracked for multiple days at high resolution (25 Hz in our case), manual classification of entire tracks is impractical. We used a supervised classification to assign behavioural states to the accelerometer data:

1. We calculated multiple metrics from the accelerometers that have been used to classify behaviours in other wildlife studies.
2. We manually classified a small subset of each individual track to nine behavioural states.
3. We trained a random forest classification model using 50% of the manually classified data.
4. We validated the performance of the random forest classification model using the withheld classifications.
5. We predicted behavioural states for the entire accelerometer dataset using the classification model.

All accelerometer analysis was performed in R, version 4.3.2 [33].

### Accelerometer-derived metrics

We calculated a range of accelerometer-derived metrics shown to be useful in characterizing bird behaviour and different modes of flight [40, 43, 45, 46] including surge acceleration, sway acceleration, heave acceleration, dynamic surge acceleration, dynamic sway acceleration, dynamic heave acceleration, body pitch, body roll, vectorial static body acceleration (VeSBA), vectorial dynamic body acceleration (VeDBA), and wingbeats. Vectorial dynamic body acceleration (VeDBA) and vectorial static body acceleration (VeSBA) were calculated following [45]. VeSBA measures gravitational acceleration which should be close to 1 when the animal is stationary or making linear movements, and greater than 1 when the bird is turning [45]. VeDBA measures total dynamic movement of the animal, which can be influenced by intrinsic movements and environmental conditions [45]. All measures except wingbeats were calculated over a 1-sec moving window. Wingbeats were identified using the `findpeaks` function in the `pracma` package [47]. Potential wingbeats were first identified as peaks in heave acceleration with an amplitude >1 g where at least two successive peaks occurred within 4 Hz, these thresholds were identified throughout visual examination of heave acceleration.

### Manual behavioural classification

We visually examined plots of the accelerometer-derived metrics at different time scales (e.g. 15 sec, 1 min, 5 min, 1 hr) to identify consistent patterns that were likely associated with nine general behaviours: three behaviours in the burrow, three behaviours associated with swimming, and three flight behaviours. From visual examination, acceleration in the heave axis, pitch, roll, VeSBA, VeDBA, and temperature were most useful in visually identifying characteristic behaviours. Table 2 provides a description of the nine behaviour classes. We randomly selected 400 15-sec segments from each bird tracked with an accelerometer (n=4400 segments, representing 0.8% of all segments). Each segment was manually classified to one of nine possible behaviours by a single observer (AP). Segments were each classified twice in random order. Segments that were not classified consistently to the same behaviour were reviewed a third time to determine a final classification.

**Table 2:** Description of the movement characteristics used to manually classify training segments to nine behaviour classes. Final column shows the number of randomly selected segments classified to each behaviours. Although 4400 random segments were initially selected, 26 segments could not be reliable assigned to a behaviour class and were removed from the training dataset.

| Behaviour | Behaviour characteristics | Training segments |
| --- | --- | --- |
| Burrow - still | Consistent high temperature; low <u>VeDBA</u>; virtually no variation in surge, sway, and heave axes | 1350 |
| Burrow - stirring | Consistent high temperature; low <u>VeDBA</u> with max <u>VeDBA</u> < 1 g; small amplitude variations in surge, sway, and heave axes | 715 |
| Burrow - active | Consistent high temperature; max <u>VeDBA</u> > 1 g; sustained variation in <u>VeDBA</u>, surge, sway, and heave axes; changes in pitch and roll greater than 10° | 338 |
| Flying - non-flapping | Consistent fluctuations in <u>VeSBA</u> with peaks > 1.5 g; consistent low frequency fluctuations in heave acceleration with peaks > 2 g; absence of wing beats | 102 |
| Flying - flap-glide | Consistent fluctuations in <u>VeSBA</u> with peaks > 1.5 g; consistent fluctuations in heave acceleration with peaks > 2 g; at least one bout of 3 successive wing beats | 540 |
| Flying - intensive | Fluctuations in <u>VeSBA</u> with peaks > 1.5 g; fluctuations in heave acceleration with peaks > 2 g; sustained bouts of wing beats > 4 sec, unusually high amplitude wingbeats (>2 g); irregular variation in pitch or roll | 70 |
| Water - inactive | Low mean and peak <u>VeDBA</u>; random low amplitude variation in <u>VeDBA</u>, surge, sway, and heave acceleration; minimal variation in <u>VeSBA</u> | 88 |
| Water - active | Mean <u>VeDBA</u> > 0.1 g and < 1 g; non-random variation in surge, sway, and heave acceleration; fluctuations in pitch or roll >10° and <50°; minimal variation in <u>VeSBA</u> | 130 |
| Water - intensive | Low variation in <u>VeSBA</u>, peak <u>VeSBA</u> <1.5 g; high mean <u>VeDBA</u> > 1g, peak <u>VeDBA</u> > 2g; large variation in pitch/roll > 50°; intense non-random variation in surge, sway, and heave acceleration | 116 |

### Random forest model training, evaluation and behavioural prediction

Manually classified data were split into training (50%) and validation (50%) data. For each 15-sec segment in the tracking data, we calculated summary statistics of the 12 metrics described in Section “Accelerometer-derived metrics”. Summary statistics included mean, inter-quartile range (IQR), 10th quantile, 90th quantile, and sum (only applied to wing-beats); a detailed explanation of how each metric was summarized is provided in Supplementary Information Table S1. This resulted in 36 potential predictor variables for behavioural classification.

We used the training data to fit a classification algorithm for the nine behavioural states. Behavioural classification was conducted using a random forest classification model with the `ranger` package (version 0.16.0, [48]). Model training and variable selection were done using the `caret` package (version 6.0-94, [49]). The `ranger` model has three parameters (minimum node size, `mtry`, and `extratrees`) that need to be optimised. This model tuning was done using repeated k-fold cross-validation with 5 repeats of 10 folds, where within each fold, 50% of the data were used in model training and 50% were withheld for model testing. Behavioural classes were up-sampled (e.g., observations of each behaviour were resampled with replacement) to ensure even sample sizes across all behaviours within the training dataset. Model tuning parameters were selected using the random grid search option with 18 different combinations of values for the hyperparameters of `min.node.size`, number of variables used at each node (`mtry`), and splitting rule (`gini` or `extratrees`). We used Recursive Feature Elimination (RFE) to reduce the number of predictor variables in the model, using the `rfe` function in the `caret` package [49]. This algorithm fit a full model with all possible predictors, then recursively removed predictors based on the variable importance ranking and refits the reduced models. Kappa was compared across all the models; the variables included in the simplest model within 1% of the highest accuracy were used in the final model. The final model was fit using 14 predictor variables (Supplementary Information Table S1) with `min.node.size` = 3, `mtry` = 11, and split rule = `extratrees`.

We used the final random forest model to predict behavioural classifications for all segments. The withheld validation segments were used to estimate the accuracy of the final model on independent data. We assessed overall model performance based on the overall classification accuracy and Kappa. We explored the behaviour-specific classification accuracy by looking at balanced accuracy for each of the nine behaviours and for the three main behavioural modes combined (burrow, water, flying). We looked at the variable importance to determine which accelerometer measures contributed the most to classification accuracy.

### Daily activity patterns

We calculated the proportion of time spent in the six at-sea behaviours over 1-hour intervals (from 8 birds, hereafter accelerometer-derived behaviours) and the proportion of time spent in the two at-sea states over 4-hour intervals (from 7 birds, hereafter GPS-derived behaviours) and estimated the sun angle (in radians) for each time step using the coordinates of the colony [50]. For each behaviour, we fit a zero-inflated beta generalized linear mixed model with sun angle as a fixed effect, random slope and intercepts for bird identity, and a first-order autoregressive correlation structure [51]. We generated posterior predictive plots to check for systematic discrepancies between model output and the observed data [52].

## Results

Data were obtained from 8 of 9 Axy5 deployments (89%), 3 of 5 AxyTrek deployments (60%), and 7 of 11 PathTrack deployments (63%). Trip duration from the Axy5 and AxyTrek devices averaged 8.2 d and 8.5 d, respectively although only three of the devices (all Axy5 units) recorded a complete foraging trip (Table 1). Three of the petrels with Axy5 devices remained in their burrow throughout the deployment. Biologgers with GPS sensors recorded eight complete foraging trips (all from PathTrack units; Figure 1) and three incomplete trips (AxyTrek units).

**Figure 1:**
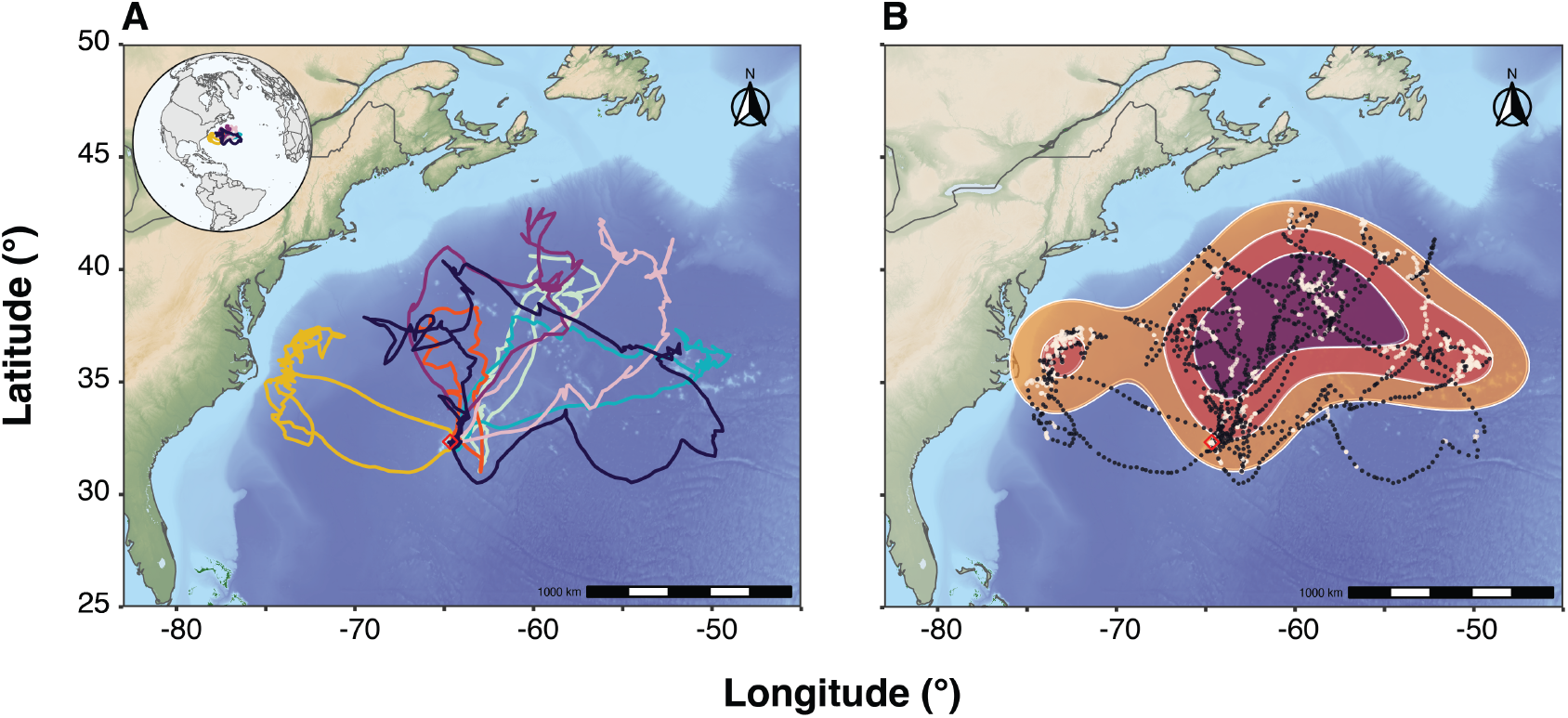
A) Foraging tracks for seven Bermuda petrels tracked from Nonsuch Island, Bermuda during incubation in 2023. Tracks show seven complete foraging trips recorded using PathTrack Nano-Fix GPS units; B) UD 95,75,50 and 25% calculated using only “search” classified location (through HMM).

### Foraging trips and at-sea distribution

During the study period, we recorded 7 complete incubating foraging trips from petrels tracked with GPS loggers (Path-track devices; Figure 1A). The mean (±SD) duration of the trips was 10.3±2.9 d with an average maximum distance travelled of 1207±305 km (Table 1). The foraging distribution (locations classified as “search” by HMM) covered an area of 1,939,342 km^2^ (UD 75%). Two main core areas (UD 25%, 438,609 km^2^) were identified in the deep waters of the western North Atlantic (Figure 1B).

### Diving behaviour

We found no clear evidence that petrels engaged in diving activity. After applying the ZOC, less than 0.001% of observations from 8 petrels had depths >0.1 m and the deepest depth recorded was 1.6 m (Figure 2).

**Figure 2:**
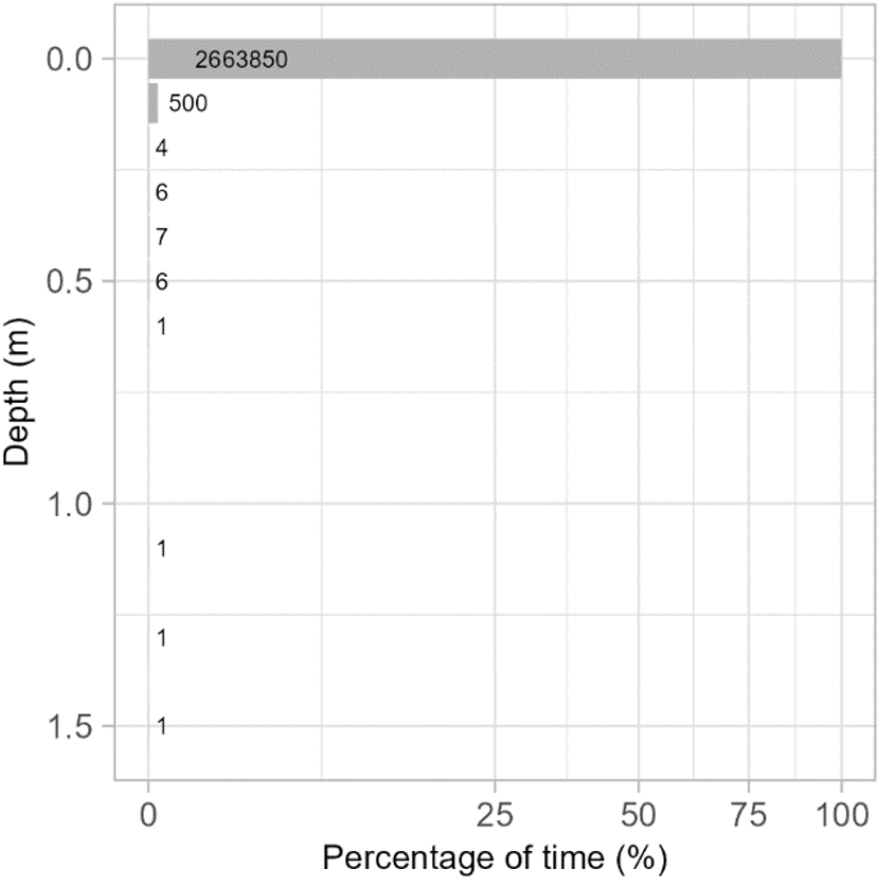
Percentage of depth measurements within 0.1 m depth classes, pooled across all foraging trips. Numbers to the right of bars represent the number of depth measurements in that depth class. The x-axis is square-root transformed to improve visualization of the range of values in the plot.

### Accelerometer behaviour classification

Overall, the random forest model predicted behaviours with 87.1% accuracy (CI: 85.7–88.5) with a Kappa of 83.9%. Most individual behaviours had balanced classification accuracy between 85.6% and 96.5%, except for the “flying-intensive” behaviour (61.7%, Table 3). When behaviours were further collapsed into three main activity modes of burrow, water, and flying, model accuracy increased to 98.7% (CI: 98.2– 99.1), with a Kappa of 97.7% and balanced accuracy for individual classes between 96.3% and 99.6%. This shows that most misclassifications occurred within these three main behavioural modes. The five most important variables in the classification were mean VeDBA (100%), the 90th quartile of VeDBA (82.3%), mean dynamic sway acceleration (78.8%), mean dynamic heave acceleration (76.1%) and the IQR of heave acceleration (74.4%, Supplementary Information Figure S1).

**Table 3:** Behaviour specific performance metrics for the random forest model predicting Bermuda Petrel behaviour from accelerometer data. Left-hand columns show results for detailed behavioural classes and right-hand columns show results for general behavioural modes.

| Behaviour | Precision | Recall | Accuracy | Mode | Precision | Recall | Accuracy |
| --- | --- | --- | --- | --- | --- | --- | --- |
| Water - inactive | 0.875 | 0.714 | 0.856 | Water | 0.936 | 0.931 | 0.963 |
| Water - active | 0.736 | 0.877 | 0.933 |  |  |  |  |
| Water - intensive | 0.867 | 0.788 | 0.892 |  |  |  |  |
| Flying - non-flapping | 0.920 | 0.955 | 0.965 | Flying | 0.982 | 0.989 | 0.989 |
| Flying - flap-glide | 0.868 | 0.890 | 0.936 |  |  |  |  |
| Flying - intensive | 0.529 | 0.237 | 0.617 |  |  |  |  |
| Burrow - still | 0.928 | 0.925 | 0.947 | Burrow | 0.998 | 0.994 | 0.996 |
| Burrow - stirring | 0.771 | 0.762 | 0.860 |  |  |  |  |
| Burrow - active | 0.800 | 0.804 | 0.894 |  |  |  |  |

#### Burrow behaviours

Burrow behaviours had low measures of activity, relative to the other behaviour classes, across all accelerometer metrics and relatively high mean temperatures (Supplementary Information Figure S2). Burrow-still had the lowest values across all measures of acceleration. Burrow-stirring had higher activity measures than burrow-still, but values were lower than for other behaviours. The burrow-active behaviour had similar values for mean VeDBA and mean dynamic surge compared to the water-inactive behaviour; however, burrow-active had lower IQR heave acceleration and lower IQR VeSBA (Supplementary Information Figure S2).

#### Water behaviours

The three water behaviours were distinguished from the three flying behaviours by lower IQR in both the heave axis and VeSBA (Supplementary Information Figure S2). Water-inactive had very low VeDBA (Supplementary Information Figure S3) and likely represented periods of resting on the water when movement was coming from the movement of the water rather than the motion of the bird. Water-active had higher activity overall than water-inactive but still had low VeDBA compared to other at-sea behaviours (Supplementary Information Figure S2). The water-active class likely represents periods of minimal activity, for example, when the bird was actively maintaining its position in the current through swimming. Water-active could include passive foraging movements, like picking at prey at the surface, but not movements consistent with active pursuit of prey or competition with other birds. Water-intensive had the highest activity measure of all the water behaviours, especially pronounced in mean VeDBA, wingbeats, and mean dynamic surge. High dynamic surge was likely associated with repeated dipping of the head into or towards the water, potentially to catch prey or as part of bathing/preening (Supplementary Information Figure S3). The occurrence of sporadic wingbeats could also have been associated with preening or short hops across the water.

#### Flight behaviours

The three flight behaviours were all characterised by high values of mean VeDBA, IQR heave, and IQR VeSBA (Supplementary Information Figure S2). Flap-glide flight and non-flapping flight had similar values for mean VeDBA, but flap-glide had higher numbers of wing-beats and lower IQR VeSBA. The large variation in VeSBA during non-flapping flight is consistent with petrels using dynamic soaring to reduce energetic costs under favourable wind conditions and using bouts of flapping to power flight when wind conditions are unfavourable to dynamic soaring (Supplementary Information Figure S4). Intensive flight had more wingbeats per second than flap-glide flight, as well as more dynamic acceleration in the surge axis. Higher wing-beats could be associated with take-offs and landings or other aerial manoeuvring. Higher dynamic heave acceleration, similar to water-intensive, could indicate dipping or plunging towards the water. The flying-intensive behaviour had much lower accuracy against the validation data than the other behaviour classes, as it likely reflects a range of short-duration behaviours that share the characteristics of unusually high wingbeats and high IQR in the VeSBA. As *Pterodroma* petrels are known to forage in flight [20, 21], it is plausible that all three flight behaviours could include foraging.

### Activity budgets

While on foraging trips, petrels spent more than three-quarters of their time in flight (Figure 3 and Figure 4), primarily in non-flapping flight (52.9 ± 14.5%, range: 26.8– 67.1%) followed by flapping flight (27.6 ± 11.2%, range:12.7–46.1%) (Figure 3). Across all birds, only a small proportion of time was classified as flying-intensive (0.9% ± 0.6; 0.2–1.7%). While on the water, petrels spent similar proportions of time active (9.2 ± 1.5%, range: 7.3–10.8%) compared to inactive (6.2 ± 4.7%, range: 0.14–11.9%), but less time in the behaviour classified as water-intensive (4.5 ± 1.5%,range: 2.3–6.4%) than in the other two water behaviours.

**Figure 3:**
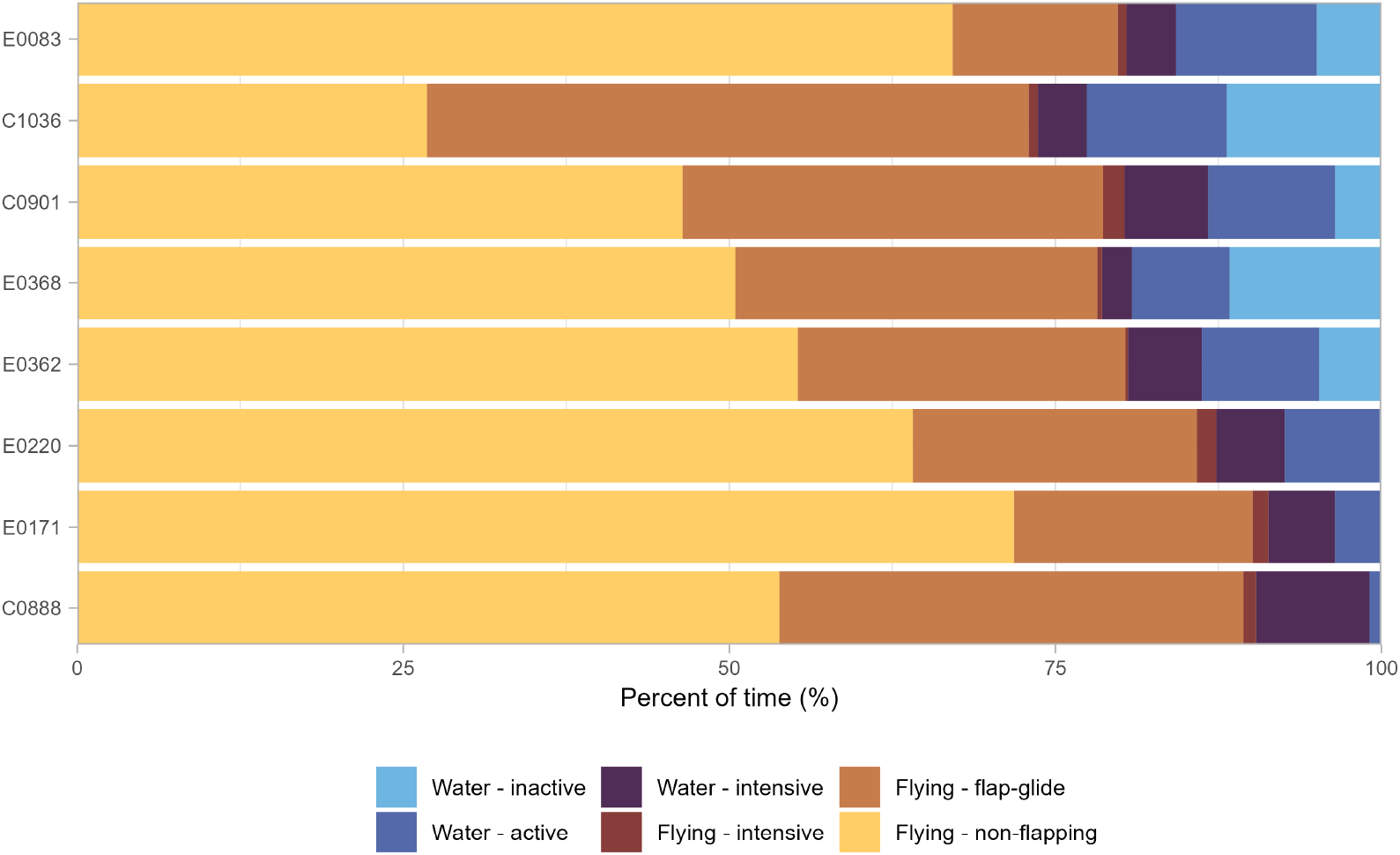
Activity budgets of the 8 Bermuda petrels tracked from Nonsuch Island with accelerometers in 2023. Colours indicate the six behaviours classified using a random forest model, individual bird identities are indicated on the y-axis.

**Figure 4:**
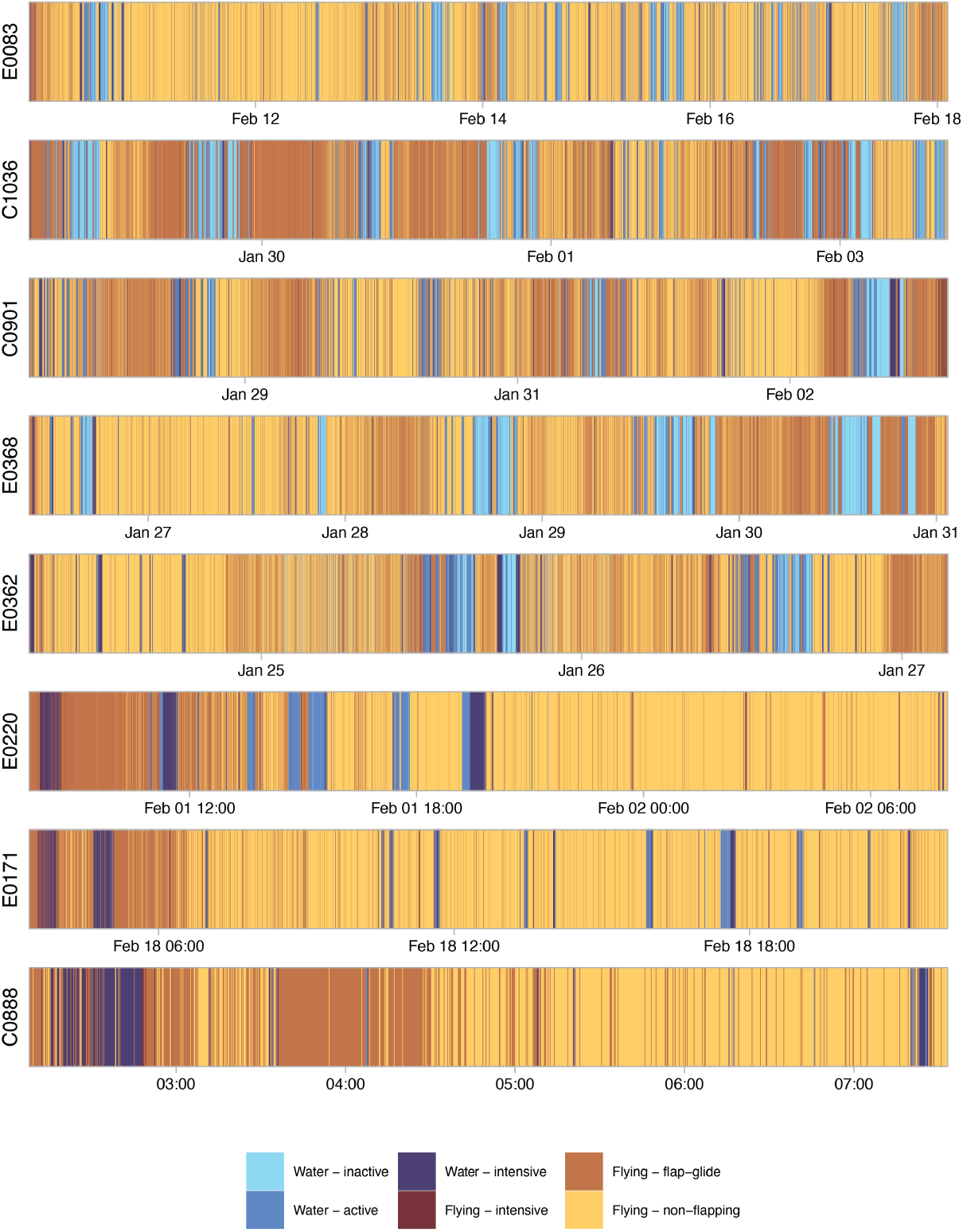
Sequence of classified behaviours within deployments of accelerometers on Bermuda petrels. Colours indicate the six behaviours classified using a random forest model, individual bird identities are indicated on the y-axis. Note that dates are different along the x-axis for each bird because deployments were not concurrent.

### Daily activity patterns

Sun angle had an effect on the proportion of time birds spent in all accelerometer-derived behaviours during foraging trips, except flying-intensive (Figure 5A-F and Table 4). Flying-non-flapping was the most common behaviour under all conditions. Both flying-non-flapping and flap-glide behaviours had a high probability of occurring at all sun angles, but the probability of occurrence declined with positive sun angles (Figure 5E-F and Table 4). The proportion of time spent in flying-flap-glide behaviour had a strong negative association with sun angle (Figure 5E). All three behaviours on the water were positively associated with solar angle (Figure 5A-C), increasing both in occurrence and proportion of time spent in the behaviour (Table 3). Interestingly, both the water-active and water-intensive behaviours had a higher probability of occurring during the day than the water-inactive behaviour (Figure 5B-C), but overall birds spent a higher proportion of their total time in water-inactive compared with water-active behaviour. This likely reflects that the resting intervals occurred more sporadically but lasted longer than other surface activities. We found no relationship between sun angle and flying-intensive, which was the most uncommon behaviour under all conditions (Figure 5D). Collectively, the results demonstrate that Bermuda petrels are primarily aerial during the night and only spend significant time on the water during the day.

**Table 4:** Model coefficients for zero-inflated beta generalized linear mixed model predicting the effect of sun angle on the probability of Bermuda petrels engaging in different behaviours during foraging trips (see Fig. 5). Values are parameter estimates ± standard errors on the logit scale.

| **Accelerometer-derived behaviours** |  |  |  |  |
| --- | --- | --- | --- | --- |
| *Behaviour* | Zero-inflated model |  | Conditional model |  |
|  | Intercept | Sun angle | Intercept | Sun angle |
| Water - inactive | 0.72 ± 0.09 *** | -1.26 ± 0.14 *** | -1.40 ± 0.14 *** | 0.57 ± 0.16 *** |
| Water - active | -1.3 ± 0.10 *** | -1.2 ± 0.14 *** | -1.99 ± 0.07 *** | 0.82 ± 0.09 *** |
| Water - intensive | -1.66 ± 0.11 *** | -0.81 ± 0.15 *** | -2.84 ± 0.07 *** | 0.14 ± 0.05 ** |
| Flying - intensive | -0.05 ± 0.08 | -0.003 ± 0.11 | -4.15 ± 0.06 *** | -0.12 ± 0.06 |
| Flying - flap-glide | -3.7 ± 0.26 *** | 1.2 ± 0.45 ** | -1.53 ± 0.27 *** | -0.92 ± 0.17 *** |
| Flying - non-flapping | -2.9 ± 0.18 *** | 1.0 ± 0.31 *** | 0.003 ± 0.35 | -0.39 ± 0.19 |
| **GPS-derived behaviours** |  |  |  |  |
| Search | -0.43 ± 0.11 *** | -0.74 ± 0.16 *** | 1.39 ± 0.12 *** | -0.09 ± 0.12 |
| Transit | -0.74 ± 0.11 *** | 0.25 ± 0.17 | 1.49 ± 0.10 *** | -0.36 ± 0.11 *** |

**Figure 5:**
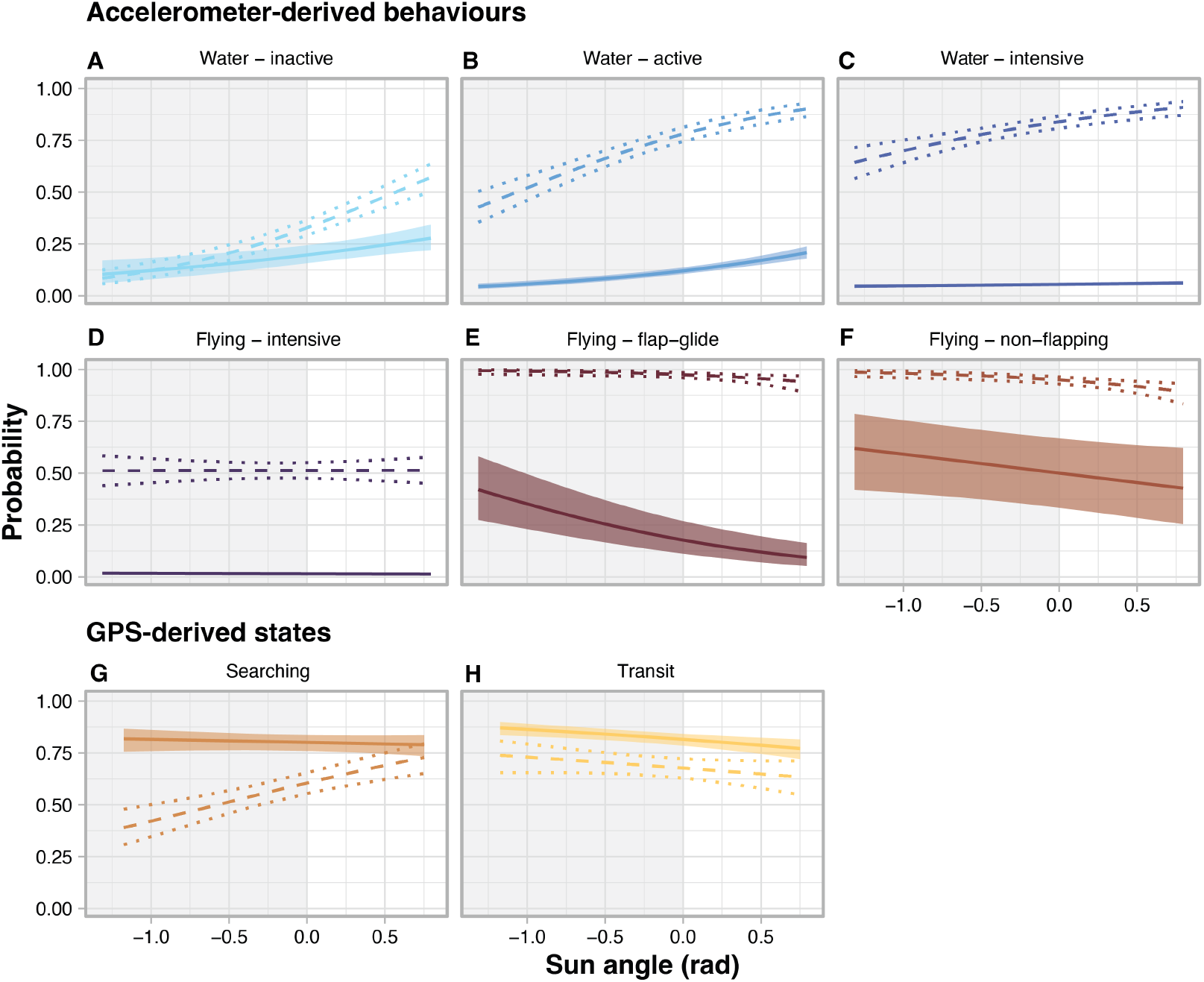
Predicted effect of sun angle on the time spent in different behaviours during trips by incubating Bermuda petrels. Models were run using behavioural modes derived from accelerometer (A-F) and GPS data (G-H). Dashed lines show the probability that each behaviour will occur and dotted lines are 95% confidence intervals. Solid lines show the proportion of time petrels were predicted to engage in each behaviour conditional on the probability that the behaviour occurs (shaded areas are 95% confidence intervals).

The probability of occurrence of the GPS-derived behavioural state “search” increased with the sun elevation (i.e., increased during the day; Figure 5G, Table 4), but the proportion of time spent searching, which may include resting on water as well as putative foraging, did not change between night and day (Figure 5G, Table 4) and was consistently high (75–80%). In comparison, the probability of occurrence and the proportion of time spent in “transit” decreased slightly during the day compared to night, but still remained relatively high (62–85%) at all sun elevations (Figure 5H, Table 4).

## Discussion

This study strongly suggests that Bermuda petrels are predominantly nocturnal surface feeders, rarely engaging in shallow diving during their extensive foraging trips. On average, petrels spent 10 consecutive days at sea, foraging thousands of kilometers from the colony along the Gulf Stream [18, 30]. This highlights their reliance on vast oceanic areas for successful foraging, a behavior also observed in other *Pterodroma* petrels [15, 53, 54]. The marine area in which the “search” activity occurred (Figure 1B) overlapped significantly with that identified for birds tracked in previous years [18, 30]; this spatial consistency in foraging areas, and the similarity observed in foraging trip metrics (duration and distance) across studies, suggests fine-scale at-sea behaviors and time budgets reported herein are representative of the species’ general behavioral patterns.

Our TDR data revealed that 99.99% of the depth measurements for Bermuda petrels were less than 0.1 m (Figure 2). In contrast to many other Procellariiformes [12, 25], which are capable of prolonged and deep dives [see references in 27, 55], *Pterodroma* petrels in general appear to not rely on dive foraging. This is likely due to the morphological and physiological adaptations that optimize their flight capabili-ties [15, 26]. Although the majority of *Pterodroma* petrels perform shallow dives, rarely exceeding two meters in depth, there are exceptions, including the Cook’s petrel and the Grey-faced petrel, which can dive to depths of approximately 27 and 23 m, respectively [25, 56]. For the Bermuda petrel feeding at the surface, our results suggest that the capture of mesopelagic and bathypelagic prey must occur exclusively when these organisms are available in the very uppermost layer of the water surface.

The accelerometer data revealed that petrels spent 80% of their time at sea in flight, which supports the hypothesis that covering long distances increases the probability of encountering prey along the route [15]. The flying-non-flapping was the most common behaviour under all conditions (mean 53%). These results support the flight strategy expected by a dynamic soaring species and are similar to the flight time budget reported for the Murphy’s petrel (95% time in flight) during incubation [57]. Such a flight strategy allows petrels and related species, including albatrosses and shearwaters, to travel for days without resting while minimizing their energy expenditure [58, 59]. Although we did not include environmental variables in our analysis, as most at-sea activity data came from biologgers without GPS sensors, previous works have shown that some petrel species select favourable wind directions to support their extremely long-ranging movements and enhance foraging success [15, 18, 60]. In general, Bermuda petrels were less active and spent more time on the water during the day compared to night—all three water behaviours were more likely, in terms of both occurrence and proportion of time, at higher sun elevations (Figure 5). A similar pattern of diurnal resting activity has been observed in Stejneger’s petrels (*Pterodroma longirostris*, 61), Trindade petrels (*Pterodroma arminjoniana*, 62), and Desertas petrels (*Pterodroma deserta*, 63), although no such pattern has been observed in two Pacific gadfly petrels, the Chatham petrel and the Murphy’s petrel, which tend to land on the water in similar proportions during foraging trips in both daylight and darkness [53, 57, 64]. The water-intensive behaviour was the least frequent of the water behaviours, characterised by bursts of “hyperactivity,” which may suggest scavenging and seizing dead and/or live prey on the water surface. Scavenging is the most likely feeding mode to explain the occurrence of very large squids in petrels’ stomach content [10, 12, 20] or big chunks of fishes found in Bermuda petrel nests (Campioni L., personal observations). However, without external validation it is impossible to be certain that this water-intensive behaviour reflects a surface foraging activity.

How birds access deep-sea prey or which prey groups are available, particularly during daylight, to specialized mesopelagic predators with limited diving ability, such as the Bermuda petrel, is not well understood. Several mesopelagic fishes, such as lanternfish, have epipelagic larvae which can be found in surface waters between 0–20 m depth [65]. Additionally, many squid species, including some preyed upon by the Bermuda petrel [18], exhibit a planktonic embryo stage (*Stigmatoteuthis hoylei* and *Histioteuthis corona* in www.sealifebase.ca), thus available to surface-feeding predators. However, the life cycle and ecology of these prey species do not fully explain the presence of fish bones and remains of adult mesopelagic fishes found in diet analyses of *Pterodroma* petrels (Silva M., unpublished data). The interaction between prey ecology and oceanographic features [66] may provide further insight into this complex predator-prey dynamic, but more research is needed to clarify these relationships.

In contrast to what we observed for water behaviours, flight behaviours increased both in frequency of occurrence and proportion of time predicted to be in flight, with negative sun elevation values. This indicates that petrels were particularly active “on the wing” at night, highlighting their pronounced nocturnality. All three flight behaviours could include the previously described “aerial dipping” of *Pterodroma* petrels [20, 21], which would then explain how the birds access the deep-sea prey that perform diel vertical migration [13].

Furthermore, nocturnality changes with breeding stage and is influenced by moon illumination [57, 64, 67, 68]. For example, the flight activity of the sympatric Zino’s and Desertas petrels, during their non-breeding season, peaked on nights with the most moonlight [69]. However, the effect of moonlight on petrel activity seems to be pronounced during the non-breeding period when birds are not constrained by their breeding duties [69].

Importantly, within and between species differences in activity budgets can occur with different tracking technologies (e.g., geolocator, GPS, accelerometer) and analytical methods used to infer behaviours. Our study highlights this issue, as the accelerometers provided a more detailed discrimination of petrel activities compared to the GPS-only devices, revealing a marked nocturnality that had previously gone undetected in GPS-based studies [18]. Overall, our results and those of studies utilizing geolocators with wet-dry sensors provide robust empirical evidence of varying degrees of nocturnality across gadfly petrels [57, 63, 70].

This study provides important new insights into the foraging behavior of an endangered species, with significant implications for its at-sea conservation. The Bermuda petrel is a highly active species, with individuals spending 80% of their time in flight while foraging over vast oceanic areas. As such, large-scale environmental changes, such as increasing SST and marine heatwaves [71], alteration in the Gulf Stream, or changes in wind regimes [4], may shift the spatio-temporal availability of food resources and affect the energetic trade-offs for this aerial predator that likely performs at the limit of its flying capacity. In addition, their pronounced nocturnal behaviour may increase their vulnerability to offshore lights that disorient birds [72], such as those associated with oil and gas infrastructure [30] and with shipping and commercial fishing vessels operating in the western North Atlantic. Further studies of their fine-scale movements across the annual cycle and over multiple years will improve our ability to protect this species at sea.

## Acknowledgments

We are grateful to J.-P. Rouja and L. Thorne for their valuable help in the field. This study was funded by Environment Climate Change Canada (ECCC). We are also grateful for the financial support through national funds to MARE (UIDB/04292/2020 and UIDP/04292/2020) by FCT, Portu-gal.

