## Supplementary material for "Fine-scale movement data reveal surface foraging and nocturnal flight activity in the endangered Bermuda petrel": Table S, Fig. S

### Table of Contents:


| | |
|---|---|
| **Table S1** | Page 2 |
| **Figure S1** | Page 3 |
| **Figure S2** | Page 4 |
| **Figure S3** | Page 5 |
| **Figure S4** | Page 6 |

**Table S1.** Description of the 36 accelerometer derived statistics used to summarize behaviour characteristics within each 15-sec segment of tracks. The last column indicates which variables were included in the final model after recursive feature elimination.

| Accelerometer measure | Summary statistic | Included in final model |
|---|---|---|
| Temperature (C) | Mean | Y |
| Surge (g) | Mean | |
| | 10th quartile | |
| | 90th quartile | |
| | Inter-quartile range | |
| Sway (g) | Mean | |
| | 10th quartile | |
| | 90th quartile | |
| | Inter-quartile range | |
| Heave (g) | Mean | |
| | 10th quartile | |
| | 90th quartile | Y |
| | Inter-quartile range | Y |
| Wing beats | Sum | Y |
| VeDBA (g) | Mean | Y |
| | 10th quartile | |
| | 90th quartile | Y |
| | Inter-quartile range | |
| VeSBA (g) | Mean | |
| | 10th quartile | |
| | 90th quartile | |
| | Inter-quartile range | Y |
| Pitch | Inter-quartile range | |
| Roll | Inter-quartile range | |
| Dynamic heave (g) | Mean | Y |
| | 10th quartile | |
| | 90th quartile | Y |
| | Inter-quartile range | |
| Dynamic sway (g) | Mean | Y |
| | 10th quartile | |
| | 90th quartile | Y |
| | Inter-quartile range | |
| Dynamic surge (g) | Mean | Y |
| | 10th quartile | Y |
| | 90th quartile | |
| | Inter-quartile range | Y |

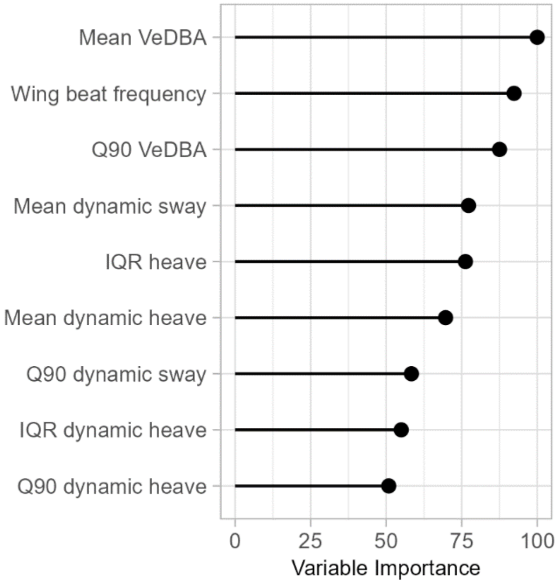


**Figure S1.** Relative variable importance for the final random forest model classifying behavioural states of Bermuda petrels from accelerometer tracks.

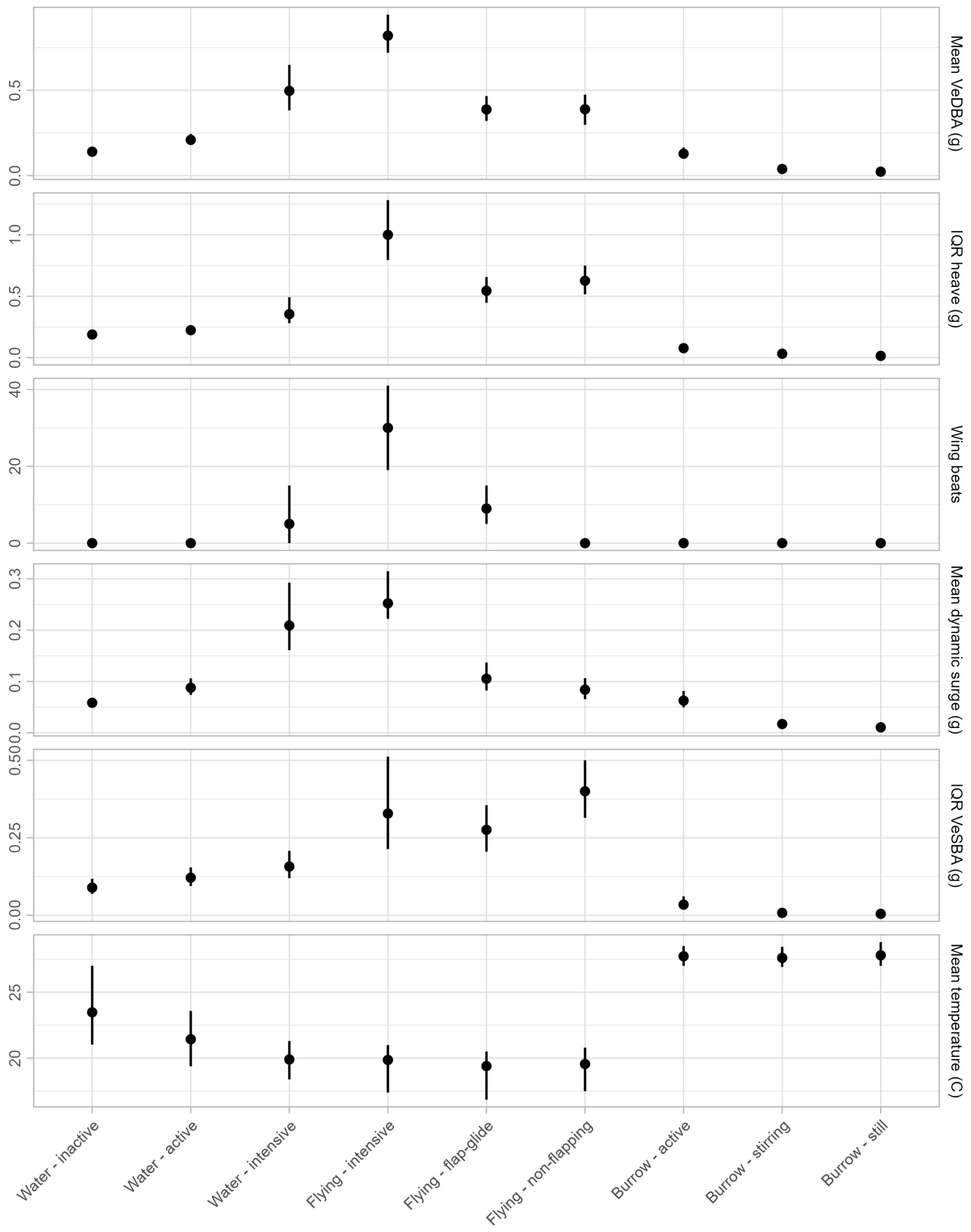


**Figure S2.** Distribution of acceleration metrics within each behaviour. Points show median value of all segments classified to a behaviour and error bars show $25^{th}$ – $75^{th}$ quantiles.

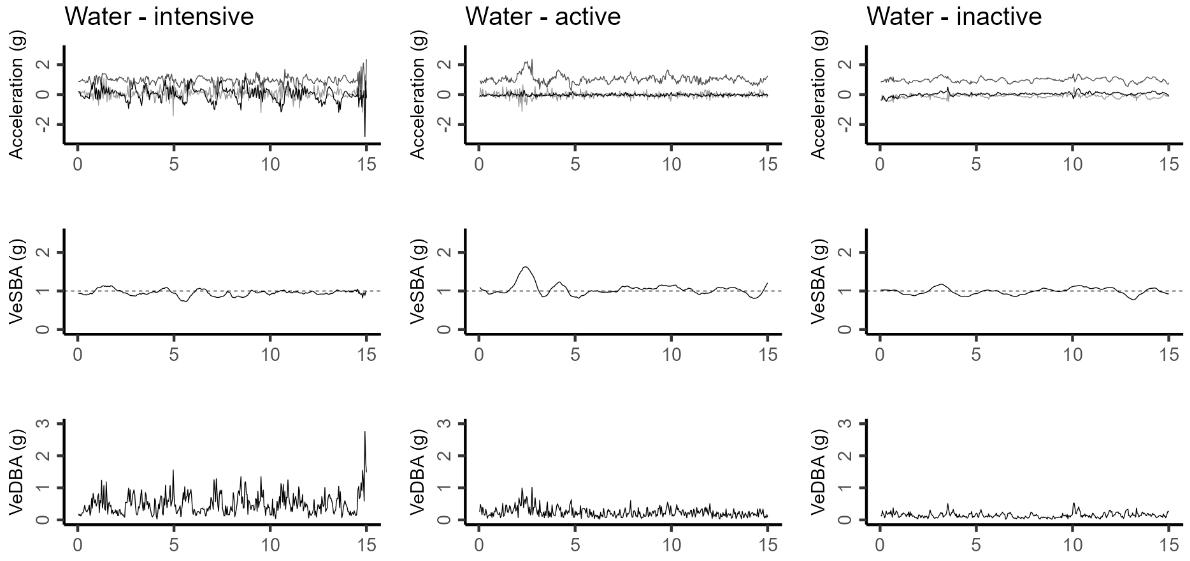


**Figure S3.** Examples of the three water modes identified from Bermuda Petrels tracked with accelerometers. Top graphs show the raw acceleration recorded surge (black), sway (light gray), and heave (medium grey) axes. Middle graphs show vectorial static body acceleration (VeSBA) which measures how much gravitation force the animal is experiencing, values of 1 g (horizontal dashed line) represent no centripetal force. Bottom graphs show vectorial dynamic body acceleration (VeDBA) which measures how much dynamic force the animal is experiencing either through internal motion or external environmental conditions.

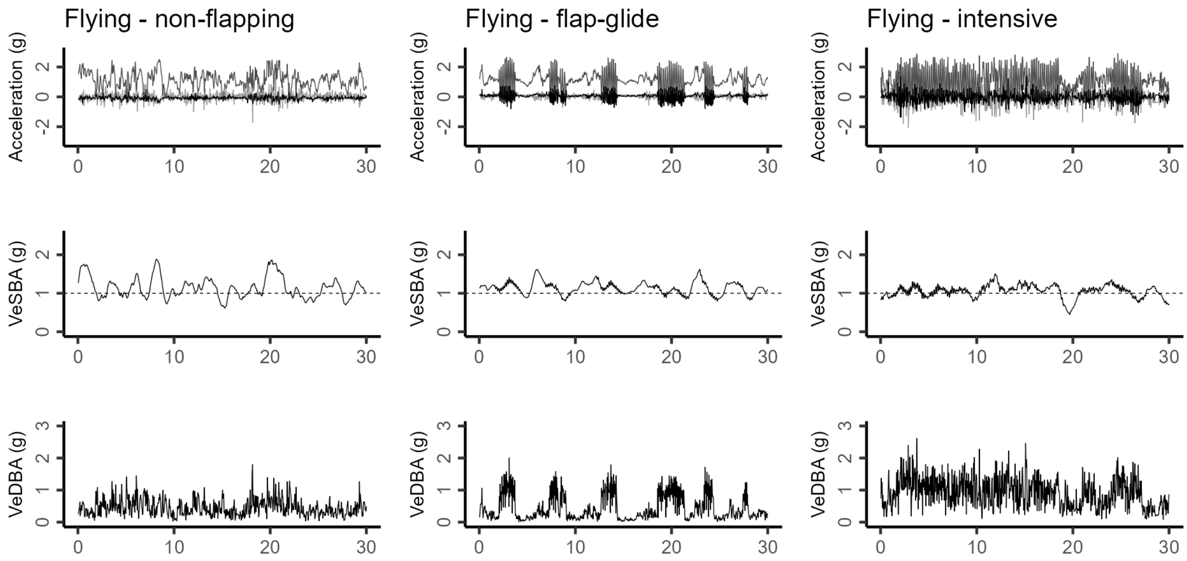


**Figure S4.** Examples of the three flight modes identified from Bermuda Petrels tracked with accelerometers. Top graphs show the raw acceleration recorded surge (black), sway (light gray), and heave (medium grey) axes. Middle graphs show vectorial static body acceleration (VeSBA) which measures how much gravitation force the animal is experiencing, values of 1g (horizontal dashed line) represent no centripetal force. Bottom graphs show vectorial dynamic body acceleration (VeDBA) which measures how much dynamic force the animal is experiencing either through internal motion or external environmental conditions.
